# Assessing the Relative Contributions of Mosaic and Regulatory Developmental Modes from Single-Cell Trajectories

**DOI:** 10.1101/2024.07.25.605053

**Authors:** Solène Song, Paul Villoutreix

## Abstract

Development is a highly complex process consisting of coordinated cell proliferation, cell differentiation and spatial organization. Classically, two ways to specify cell types during development are hypothesized : mosaic and regulative modes. In the mosaic mode, a particular cell isolated from the rest of the embryo will nevertheless give rise to cells with a fate identical to the ones expected in normal development, thus relying on lineage-inherited factors. In the regulative mode, the fate of a cell depends on its interactions with its environment, and thus relies on space-dependant factors. Both modes often coexist in the development of a given animal. We propose to quantify their respective contributions from single-cell trajectories. *C. elegans* development provides a unique opportunity to elaborate such an approach. Indeed, its invariant lineage enables the integration of spatial positions, lineage relationships and protein expression data. Using the single cell protein expression profile as a readout of the cell state, we relate the contributions of the mosaic and the regulative modes to the following measurable quantities. The contribution of the mosaic mode, or lineage-inherited contribution is quantified by the strength of the relationship between the cell-cell *lineage distance* and the cell-cell *expression distance*. Similarly, the contribution of the regulative mode, or context-dependent contribution is quantified by the strength of the relationship between the cell-cell *context distance* and the cell-cell *expression distance*. The cell-cell *context distance* measures the similarity between the spatial neighborhoods of two cells based on the gene expression profiles of their neighbours. We assess the significance of these contributions by comparing the empirical results obtained on *C. elegans* data to artificial models generated using simple rules. With these measures, we show the co-existence of mosaic and regulative modes in the development of *C. elegans*. The relative contribution of these two modes varies across the different tissues and in time. In particular, we see in the skin tissue that during early development, the mosaic mode dominates while at later stages, regulative mode dominates, suggesting a convergence of single cell trajectories. These measures are general and can be applied to other datasets that will be made available with the progress of spatial transcriptomics and lineage-tracing, paving the way for a quantitative, unbiased and perturbation-free study of fundamental concepts in developmental biology.

## Introduction

When studying embryonic development, one wants to understand how a single fertilized egg (a zygote) becomes a fully functional organism through the coordinated dynamics of myriad of cells, which will generate the desired shape and acquire the right identity. This is a highly complex process consisting of an interplay of coordinated cell proliferation, cell differentiation and spatial organization [1]. Classically, two modes of development are hypothesized: mosaic and regulative [2] (Figure 1). In mosaic development, a particular cell isolated from the rest of the embryo will still produce the same cells as in normal development [3]. In regulative development, the fate of a cell depends on its interactions with its environment. Both modes often co-exist in the development of a given animal [2]. In experimental embryology, perturbative experiments were used to assess the respective contributions of the two modes. This can be done through isolating a cell or group of cells, grafting, or bisecting embryos to study the fate of cells when placed in unusual neighbourhood. For example, Roux experiment in 1888 consisting in destroying one blastomere in the two-cell stage of frog embryo, obtaining half an embryo, illustrates mosaic development. On the contrary, Driesch (1902) dissociated the the 2-cell stage sea urchin embryo into isolated cells and some of these isolated blastomeres gave rise to a smaller but complete sea urchin embryo, illustrating regulative development, where missing neighbours can be compensated for. These emblematic experiments, although impressive, do not provide a fully interpretable picture. Indeed they provide an answer only for the specific stage in development. At later stages, when cells are more numerous, different combinations of cells removal should be tested, which would quickly become intractable. In addition, they are highly disruptive for the underlying processes and might interfere with various aspects of development. Molecular studies suggest that both modes are at play, as reviewed by Fraser and Harland [2]. However, there is a lack of a method to assess quantitatively the contributions of both modes across the full timeline of development.

**Figure 1.**
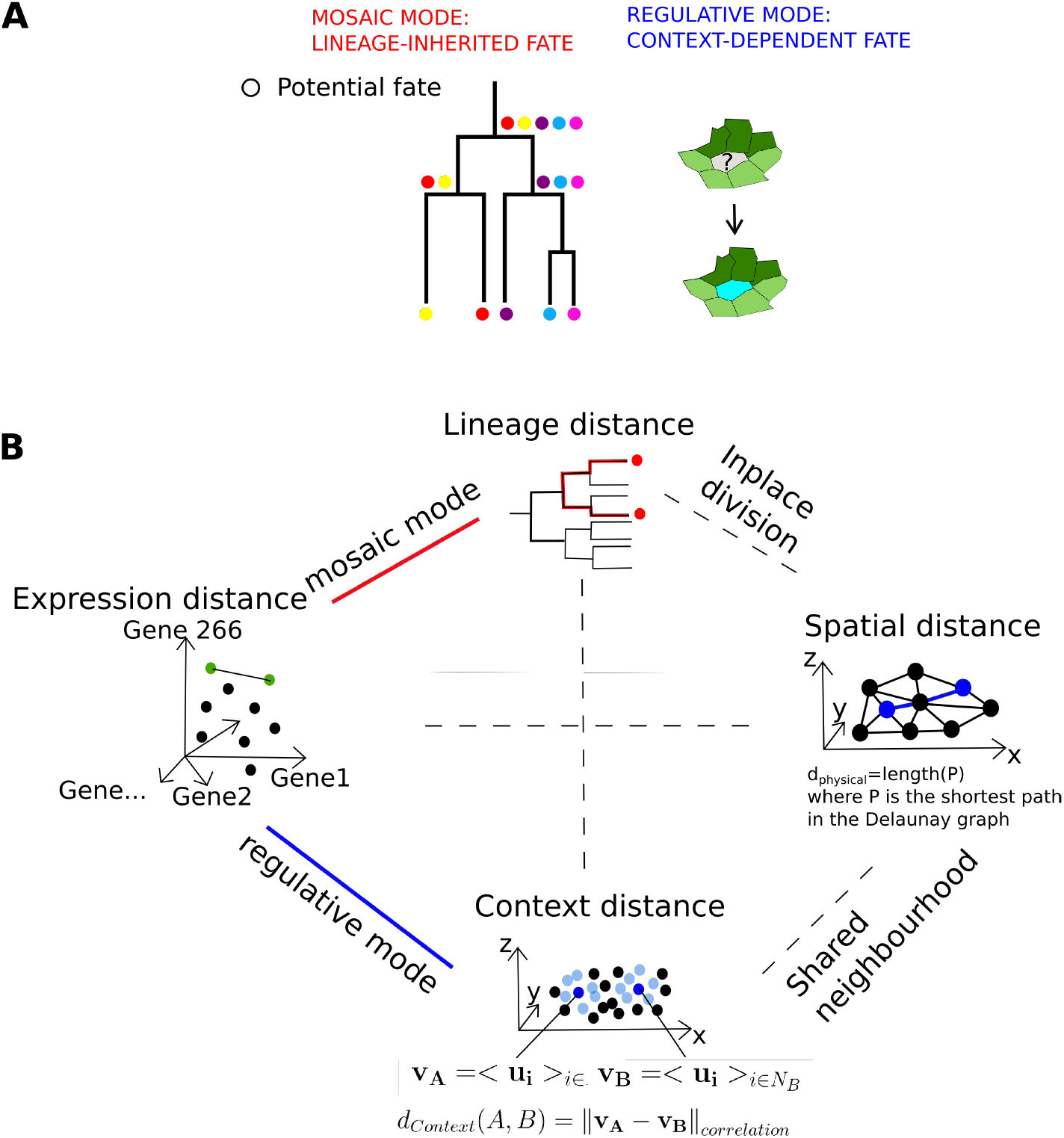
A) Schematic illustration of the distinction between mosaic mode and regulative mode in cell fate determination. In the mosaic mode, the colored dots represent potential cell fates that a cell can take, transmitted to the daughters at each division. In the regulative mode, the cell fate of the central cell is determined by the states of its neighbors represented by the two shades of green. B) Schematic illustration of the four distances used to compare individual cells, and the relations between them. The *lineage distance* is the length of the shortest path between two cells in the lineage tree taken as a graph. The *spatial distance* is the length of the shortest path in the graph of the cells physical positions. The *expression distance* is the correlation distance between two expression profiles. The *context distance* between cell A and cell B is the correlation distance between the mean expression profile of the neighbors of A and the mean expression profile of the neighbors of B.

We propose here a methodology to derive these contributions directly from measurements obtained on wild-type embryos. Our approach is made possible by recent progress in multi-modal data acquisition techniques as they enable to track cell characteristics, such as gene expression and spatial positions, along divisions at single-cell level [4, 5]. Our approach requires to have access to lineage relationship, 3D positions, and expression profiles for each cell in an embryo. The simultaneous measurement of these cell features is very challenging. Often, one aspect is inferred from others. For example, in the Morphoseq algorithm applied on the ascidian *P. mammillata* [6], the 3D positions are reconstructed from known expression profiles.

Other approaches infer cell state trajectories from the expression profiles [7], knowing that cell state trajectories are not always one-to-one with cell lineages [8]. The reconstruction of cell lineage itself can be performed from imaging data [9, 10] or barcoding techniques [11, 12]. *C. elegans*, however, enables the establishment of an integrated dataset combining lineage, 3D positions and expression profiles. Indeed, its invariant lineage development allows to combine spatial positions, lineage relationships and protein expression data. Each cell generated through development is attributed a cell identity, and has a corresponding counterpart in each embryo. The integrated dataset that we use here combines spatial positions and lineage relationships obtained by tracking nuclei [13] and protein expression dataset generated by combining expression levels mapped by imaging of 266 transcription factors [5].

*C. elegans* was often taken as the textbook example for mosaic development due to its invariant lineage [2]. Indeed, the patterns of division and differentiation are the same from an individual embryo to another. This reproducibility has led to the simplified conlusion that the developmental program must be lineage-based. However, the reproducibility of lineage also implies reproducibility of cells positioning and neighborhoods so that it is not possible to exclude the confounding explanation that cell fate is due to reproducible interactions with their neighbours [13] [14]. Laser ablation experiments [15] had shown that both modes co-exist. Later, specific mechanistic explanations were found in favour of both modes. P-granules posterior segregation that leads to asymmetric divisions that is responsible for the germ line is a clear example of lineage dependency [16] [17] [18]. The interaction of ABp blastomere with P2 via Notch signaling makes it distinguishable from ABa [19]. Another example of a regulative contribution, later in development, is the case of the left-right asymmetry of neurons development which is also due to Notch signaling [20]. Overall, both modes contribute in the development of *C. elegans* and their respective contributions have not yet been quantified.

The quality and tractability of the *C. elegans* data makes it an ideal model to develop a method to infer the contributions of mosaic and regulative modes. Indeed, it is possible to simultaneously measure *lineage distance, spatial distance* and *expression distances* for all pairs of cells that co-exist. Taking protein expression profiles as read-outs of cell state, we can reformulate the contribution of the mosaic mode as the strength of the relationship between *lineage distances* and *expression distances*. As for the contribution of the regulative mode, we define a *context distance*, which summarizes the expressions of the neighbours of the two cells of interest. In a regulative mode, two cells, which can be distant in the physical space, can have similar expression because they lie in similar neighbouring cells. These strengths are defined using a statistical comparison of stratified correlations [21] between real *C. elegans* data and a random null model. Such correlation strengths will be considered as proxys for the contributions of mosaic and regulative modes. While not to be confused with identifying causal relationships, measured correlations suggest the existence of underlying mechanisms [22].

By analyzing separately the subpopulations of cells that give rise to each of the five distinct tissues in *C. elegans*, we show that the contribution of the mosaic mode is significant in all tissues whereas the regulative mode contribution depends on time and on the tissue: it is at play earlier in Neuron and Skin than in Muscle and Pharynx, whereas it never is at play in Intestine development on the period of observation.

Besides, we highlight the existence of a population of cells in which the regulative mode is clearly dominant: that have similar fate while having similar context despite being distant in lineage and in space. In addition, our results suggest that not only proliferation, but also cell rearrangement can be responsible of the relation between the *context distance* and the *expression distance*. Last, we show an illustration of tracks of two descent lines that lead to two Skin cells, distant in lineage, distant in space, as an example of convergence which might be context driven.

## Materials and Methods

We introduce a quantification method to assess the respective contributions of the mosaic and regulative modes (*W*_*mosaic*_ and *W*_*regulative*_) in *C. elegans* development. We first introduce the datasets for *C. elegans* development. We then detail the definitions of the distances used to build the calculations of the contributions *W*_*mosaic*_ and *W*_*regulative*_. We introduce the two artificial models : the random model and the lineage-defined toy model used as comparison to the *C. elegans* dataset. We finally detail the computation of *W*_*mosaic*_ and *W*_*regulative*_, which builds on correlations between the distances previously defined and with reference to previously introduced random model. Lastly, we detail the definition of the quadrants and rearrangement rate.

### Datasets

For most cells in *C. elegans* development, we can access the following information:

- its position in the lineage tree
- its position in 3D space
- its partial protein expression profile

The lineage is published in Sulston et al. [15] We combine two other published datasets to relate each cell to it’s 3D position and protein expression profile [5, 13].

### Spatial positions

The dataset containing the spatial positions was originally published in [13], which measured the 3D positions by tracking nuclei with ubiquitously expressed mCherry. The timepoints range from t=0 to t=190, with intervals of 75 seconds. Out of the 28 embryos recorded in their dataset, we chose the embryo number 13, which had the highest number of cells. At the last time point t=190 (about 6 hrs), there are 380 cells. By this stage, the embryos have completed all but the last round of cell divisions, and have produced 762 cells, for a total number of 1341 different cells observed across the entire development.

### Expression profiles

The expression dataset was originally published in [5]. The dataset was obtained by fluorescent microscopy imaging of the protein expression levels of 266 transcription factors using protein-fusion reporters. Thanks to *C. elegans* lineage invariance, the expression profiles are unambiguously paired with the cell identities. We have the expressions of all cells including the last division, except from the first cells P0, P1 and AB, with a total of 1204 cells tracked out of the 1341 cells generated. The terminal cells come with their tissue annotation: Pharynx, Skin, Neuron, Muscle and Intestine. We propagate this terminal tissue label upstream following daughter to mother branches to identify all the cells that contribute to a given tissue. The lineage and context dependency contributions are calculated for these groups of cells separately.

### Distances

The computations for the contributions *W*_*mosaic*_ and *W*_*regulative*_ rely on the correlation between the *expression distances* and, respectively, *lineage distances* and *context distances* between pairs of cells. We detail here the definitions of these distances.

**The *lineage distance*** is the length of the shortest path between two cells in the lineage tree. The lineage tree is encoded as a network, where the nodes are the cells and the undirected edges encode the ascendent/descendent relationships. The length of the shortest path between two cells in the lineage tree is the number of edges in the shortest path from one cell to the other.

**The *expression distance*** is the correlation distance between two expression profiles using scipy.spatial.distance.correlation from Scipy library. 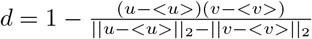

**The *spatial distance*** is the length of the shortest path in the spatial network. The spatial network is obtained by computing a Delaunay graph from the individual cell spatial positions. The cell spatial positions correspond to the center of the detected nuclei from microscopy imaging [13].

**The *context distance*** between two cells A and B is the correlation distance between the mean expression profile of the direct neighbors of A and the mean expression profile of the direct neighbors of B.

### Artificial models

We build two artificial expression models as comparison references for the *C. elegans* statistics. The first one, the random model, serves as a null model which statistics are used in the calculation of the contributions *W*_*mosaic*_ and *W*_*regulative*_. The second one, the lineage-defined model, was build to be an extreme case of mosaic development.

### Random model

The expression profiles of cells were randomly shuffled across the lineage tree. Because the spatial positions were stored using the identity in the lineage tree, the expression profiles are also randomly distributed across space.

### Lineage-defined model

Artificial expression vectors were generated so that the expression retains as much as possible the lineage tree information as an extreme case of lineage dependant development. It means that we wanted the correlation distances between two cells expression vectors to correlate very well with their *lineage distance*. We propose an approach inspired by the PROSSTT algorithm [23]. We first draw a random expression profile for the egg P0 from a uniform distribution. Then, we simulate expression change within a cell cycle by 3 steps of a random walk : *x*(*t* + 1) = *x*(*t*) + *v*(*t*), *v*(*t* + 1) = *v*(*t*) + *ϵ* where *ϵ* is drawn from a normal distribution. At each divisions, the first position of the daughters cells are made to coincide with the last expression of the mother cells. 200 gene expressions are generated this way, and the average expression over the 3 steps of random walk is taken as the measured expression of the toy model. We checked that the *expression distance* is highly correlated with the *lineage distance* as expected (*r >* 0.85).

### Calculation of the contributions *W*_*mosaic*_ and *W*_*regulative*_

An intuitive approach would simply be to calculate the Pearson correlation between *expression distances* and *lineage distance* as a proxy for the contribution of mosaic mode, and the Pearson correlation between *expression distances* and *context distances* as a proxy for the contribution of regulative mode. However, these values are both mildly high, respectively 0.49 and 0.59 (mixing all time points), and these correlations are confounded by the *spatial distances* (Supp Fig1). Thus, to assess the contributions separately, we need to perform stratified correlation. To assess the contribution of mosaic mode, we would calculate the correlation between *lineage distance* and *expression distance* among pairs that have the same *context distance*. The same goes for regulative mode, replacing inverting lineage and context. The stratified correlations obtained this way must then be compared to reference distribution to provide a statistically significant measurable value, *W*_*mosaic*_ and *W*_*regulative*_. This reference distribution comes from the random model introduced before.

### Calculation of the contribution of the mosaic mode *W*_*mosaic*_

The contribution of lineage-inherited factors *W*_*mosaic*_ characterizing the mosaic mode of development is computed for each time point. We use the Pearson correlation coefficient between *lineage distances* and *expression distances*, stratified by the *context distances* as follows.

1. We separate all the values of the *context distances* taken over time into 10 bins {*C*_1_, *C*_2_, …, *C*_10_}. For a given bin *C*_*i*_ which is represented at that time point, we compute all the *lineage distances* and the *expression distances* and derive a value of Pearson correlation 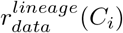, using Scipy library’s function scipy.stats.pearsonr. We obtain then 10 different values of 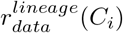.
2. We perform the same calculation in the null model at the same time point, which gives also different values of 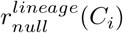.
3. We compare the distributions of 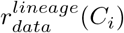 and 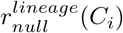 using a Mann Whitney U test (scipy.stats.manwhitneyu). The test gives us the p-value and the U statistics 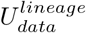 and 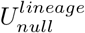 which are related to the sum of ranks of the first sample, 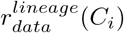, and of the second sample,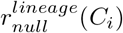.
4. From 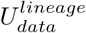 and 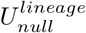 we calculate the size effect (also called rank-biserial correlation), which is how we define 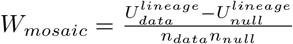 (where *n*_*data*_ and *n*_*null*_ are the number of values in the samples, here *n*_*data*_ = *n*_*null*_ = 10) which measures to what extent the stratified correlations in the real *C. elegans* data are higher than in the random null model. *W*_*mosaic*_, or effect size, is our proxy to measure the contribution of lineage-inherited factors, ie the mosaic mode.

### Calculation of the contribution of the regulative mode *W*_*regulative*_

The contribution of the space-dependent factors *W*_*regulative*_ characterizing the regulative mode of development, is computed for each time point. We use the Pearson correlation coefficient between *context distances* and protein distances, stratified by the *lineage distances* as follows.

1. For a given *lineage distance l*_*i*_ that is represented at that time point, we compute all the *context distances* and the *expression distances* and derive a value of Pearson correlation 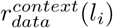, using Scipy library’s function scipy.stats.pearsonr.
2. We perform the same calculation in the null model at the same time point, which gives also different values of 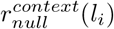.
3. We compare the distributions of 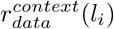 and 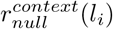 using a Mann Whitney U test (scipy.stats.manwhitneyu). The test gives us the p-value and the U statistics 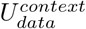 and 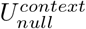 which are related to the sum of ranks of the first sample 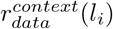, and of the second sample,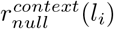.
4. From 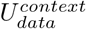 and 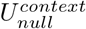 we calculate the size effect (also called rank-biserial correlation) which is how we define 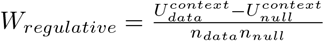 (where *n*_*data*_ and *n*_*null*_ are the number of values in the samples: the number of different values of *lineage distances* present at the given time point) which measures to what extent the stratified correlations in the real *C. elegans* data are higher than in the random null model. *W*_*regulative*_, or effect size, is our proxy to measure the contribution of the space-dependent factors, i.e. the regulative mode.

### Quadrants definition

For a given time point, each pair of cells is positioned as one data point in a 3D space, its coordinate being the *lineage distance*, the physical distance and the *context distance* between the two cells of the pair. We separate this space into 8 cubic regions (or quadrants) using the mid-point between the minimal value and the maximal value for each axis. Because the *lineage distances* and physical distances are integers, when the midpoint falls on a value that is represented in the data points, we made the choice that the data points that have exactly this value are included in the lower quadrant.

### Rearrangement rate

For each time point, for each cells, its neighbors are listed as the cells which are one edge away in the Delaunay graph. The rearrangement rate is computed by comparing the list of neighbours at time t and at time t-1 by calculating the Jaccard distance 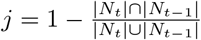

## Results

We propose a quantitative framework to assess the respective contributions of the mosaic and regulative modes in *C. elegans*. For this purpose, we build on correlation analysis at single-cell level between the *expression distance* and the *lineage distances* and between the *expression distance* and the *context distances*, respectively.

The simplest way to quantify the strength of the relationship between *expression distances* and *lineage* or *context distances* is to calculate the Pearson correlation *r*. However, one can immediately see as shown in Figure 1B that these distances are intertwined through confounding variables. If we calculate the Pearson *r* between each pairs of distances, all the *r* are mildly positive: ranging from 0.33 to 0.59 (see values in supplementary figure 1). It is possible to list a certain number of potential confounding effects. For example, because cells divide mostly in-place, it is expected that cells close in lineage, which just branched off from a common ancestor are still close in space [24]. Similarly, cells that are close physically are bound to share a number of common neighbors, and thus have a small *context distance*. Nevertheless, even if these 6 relationships between all pairs from 4 types of distances are intertwined, we can initially assume that the two, relating *expression distance* to *lineage distance* on one hand, and *expression distance* to *context distance* on the other hand, carry the core mechanistic explanations [4, 5]. Below, we use our newly introduced quantities *W*_*mosaic*_ and *W*_*regulative*_ (Methods) based on these two relationships (*expression distance* versus *lineage distance* and *expression distance* versus *context distance*) to characterize the contribution of the mosaic and regulative mode. Our results show that both modes play a role in development, depending on timing and tissue. We then highlight the effect of the context, independently of lineage or spatial relationship in convergent cell fates. In contrast, we show that the lineage dependency relies only on cells that are clonally-related. After that, we highlight that both proliferation and spatial rearrangements effects are recorded by the contributions measures *W*_*mosaic*_ and *W*_*regulative*_. Finally we provide an illustration of a pair of skin cells, distantly related in lineage, that follow different trajectories in the expression space and physical space.

### Quantification of mosaic and regulative contributions, based on stratified correlation compared to a random model, reveals diverse modes of cell specification in major *C. elegans* tissues

To assess the contribution of mosaic versus regulative modes of development, we developed two measures, for respectively lineage and context dependency contributions in time. Since these two measures are built the same way, we will explicit only one of them, the context dependency contribution. A natural way for characterizing the context dependency contribution is to evaluate the strength of the correlation between *context distances* (the factor we assess) and the *expression distances* (the read-out). We want to measure this strength independently of the *lineage distance* which is likely to be a confounding factor in this correlation. Thus we perform a stratified correlation study. At a given time point, the co-existing pairs of cells are separated according to their *lineage distance*. For each given *lineage distance* value, the correlation between the *expression distances* and the *context distances* are computed. Thus for each time point, we have several values of correlation strength, one for each *lineage distance* that exists at that time point. Similarly, we computed the correlation between *expression distances* and *lineage distances*, stratified by the *context distances*.

To evaluate if the distribution of correlation strength is high or low, we resorted to the comparison to a random model in which the expression profiles are randomly shuffled among all cells of the lineage tree. The comparison consists in calculating the Mann Whitney U test shift *W*_*regulative*_ between the *C. elegans* distribution and the null model distributions (see Methods for the definition of *W*_*regulative*_). The higher *W*_*regulative*_, the more *context distances* are correlated to *expression distances* than in the null model. These plots are shown in Figure 2 (in blue). The blue horizontal full and dotted lines mark the mean and standard deviation of the intrinsic variation of *W*_*regulative*_ of the null model (by computing the U test between all pairs of time points within the null model). The dots are filled when the p-value of the U is significant. In summary, our results indicate that there is a significant contribution of the context when there are filled blue dots above the upper blue dotted line.

**Figure 2.**
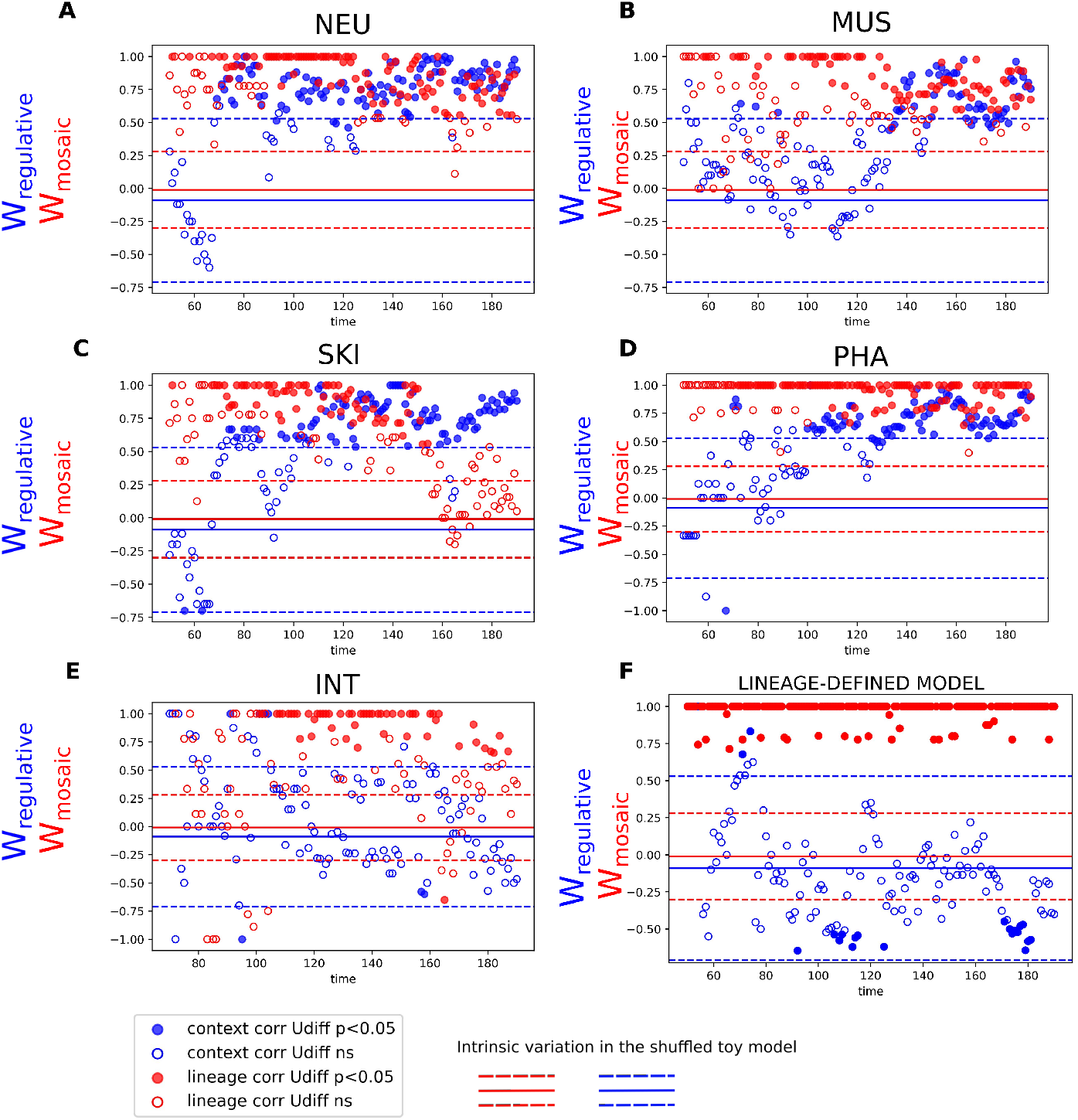
Lineage (red) and spatial context (blue) contribution to the various tissue types determination in *C. elegans* (A-E) and in a toy model (F). The lineage contribution (red) is quantified as the size effect *W*_*mosaic*_ in the statistical comparison (Mann Whitney U test) of the distributions of correlations between *expression distances* and *lineage distances* stratified by *context distances* in the given *C. elegans* tissue and the random null model. For a given time-point, the lineage contribution is considered confidently high if *W*_*mosaic*_ is filled (*p* < 0.05) and above the higher dotted red line that represent the level of intrinsic variation of the random null model. The same goes for the context contribution in blue, where the correlations are between *expression distances* and *context distances*, stratified by *lineage distances*. Lineage contribution is high in all tissues except from the end of Skin determination (C). In Neurons and Skin determination (A and C) the context contribution starts at respectively t=69 and t=71 whereas in Muscle and Pharynx (B and D), it starts later at t=136 and t=100, and it never is high in Intestine determination. In the lineage defined toy model (F), the lineage contribution *W*_*mosaic*_ is always high and the context contribution *W*_*regulation*_ never is, as expected by construction, validating the consistency of *W*_*mosaic*_ and *W*_*regulative*_ as a proxy for the lineage and context contributions.

In addition to this test, we made a toy model in which the expression profiles are artificially entirely lineage-dependent. Performing the same analysis on this lineage-determined toy model, we see, in contrast to the results with the real *C. elegans* data, that there are very few filled blue dots. This result confirms the significance of the strong correlation found in the real *C. elegans* data 2 F).

We performed this stratified correlation in comparison with the random model separately for each tissue with labels obtained from the expression dataset [5]. We consider as being part of a tissue all the cells that belong to tracks leading to the leaves that are labeled as this tissue. The context dependency contribution depends on the tissue type and on the time. Indeed, in the development of the intestine, the *context distance* is never significantly correlated with the *expression distance*. In the other 4 tissues, the context dependency is important for a period of the development. This period starts at different time points depending on the tissue. In Neurons and Skin, this period start at respectively t=69 and t=71 whereas in Muscle and Pharynx, it starts later: t=136 and t=100.

Likewise, to quantify the lineage dependency contribution, that accounts for the mosaic development, we compute the stratified correlation of *lineage distances* against *expression distances* with respect to the *context distance*. Because *context distance* is continuous, we bin the values into 10 bins. The lineage dependency persists over almost all times, except at the end of development in Skin. In the early time points, the contribution of the lineage dependency are not statistically significant due to the small number of cells. However the general trend at these early time points is that the lineage dependency contribution is beyond the intrinsic null model standard deviation line. At later times, there are statistically significant points beyond the shuffled model intrinsic standard deviation line until t=190 in all tissues except for Skin in which the lineage dependency drops from t=159 and becomes not statistically significant.

We have analyzed the relationship between the *expression distances* and *context distances*, stratifying with *lineage distances*, and the relationship between the *expression distances* and *lineage distances* stratifying with *context distances*. However, we have not ruled out the *spatial distance* as a confounding factor. In the following, we will analyze the relationship between *spatial distances* and respectively *context distances* and *lineage distances*.

### Significant effect of the context, independently of lineage or spatial relationships, suggests context led convergence effect

For each time point, we consider together the *lineage, spatial* and *context distances* for all pairs of cells that exist at that time point. Then, we classify the pairs in 8 different quadrants that each represents a configuration. The classification across the quadrants is set by the mid-distance between the minimal and maximal value of each type of distance. For instance, the blue quadrant in Figure 3 comprises the pairs of cells that are closely related in lineage, in space and in context. To characterize the relationship between *spatial* and *context distance* we are interested in the red quadrant: the pairs that share similar context while being distant in space and in lineage. This is the quadrant in which the context is entirely disentangled from the other distances. Figure 3 show that this quadrant comprises between 10 and 50% of cells along time points between t=50 and t=190. The *expression distances* of this red quadrant are higher than the clonally-related pairs of cells, but lower than the unrelated cells. The fact that cells that are context-related while being physically and genealogically distant exist and are close in expression illustrates that there is a population of cells that are apparently mainly determined by the context. In these pairs of cells, the effect of context can be isolated from the lineage and spatial effects. Comparatively, in the lineage-determined toy model, the red quadrant is represented but with a small proportion of cells, and the corresponding *expression distances* are not smaller than the unrelated cells. In the random null model, the red quadrant comprises a small proportion of cells from time t=68 (less than 7%), and has similar *expression distances* as the unrelated pairs of cells.

**Figure 3.**
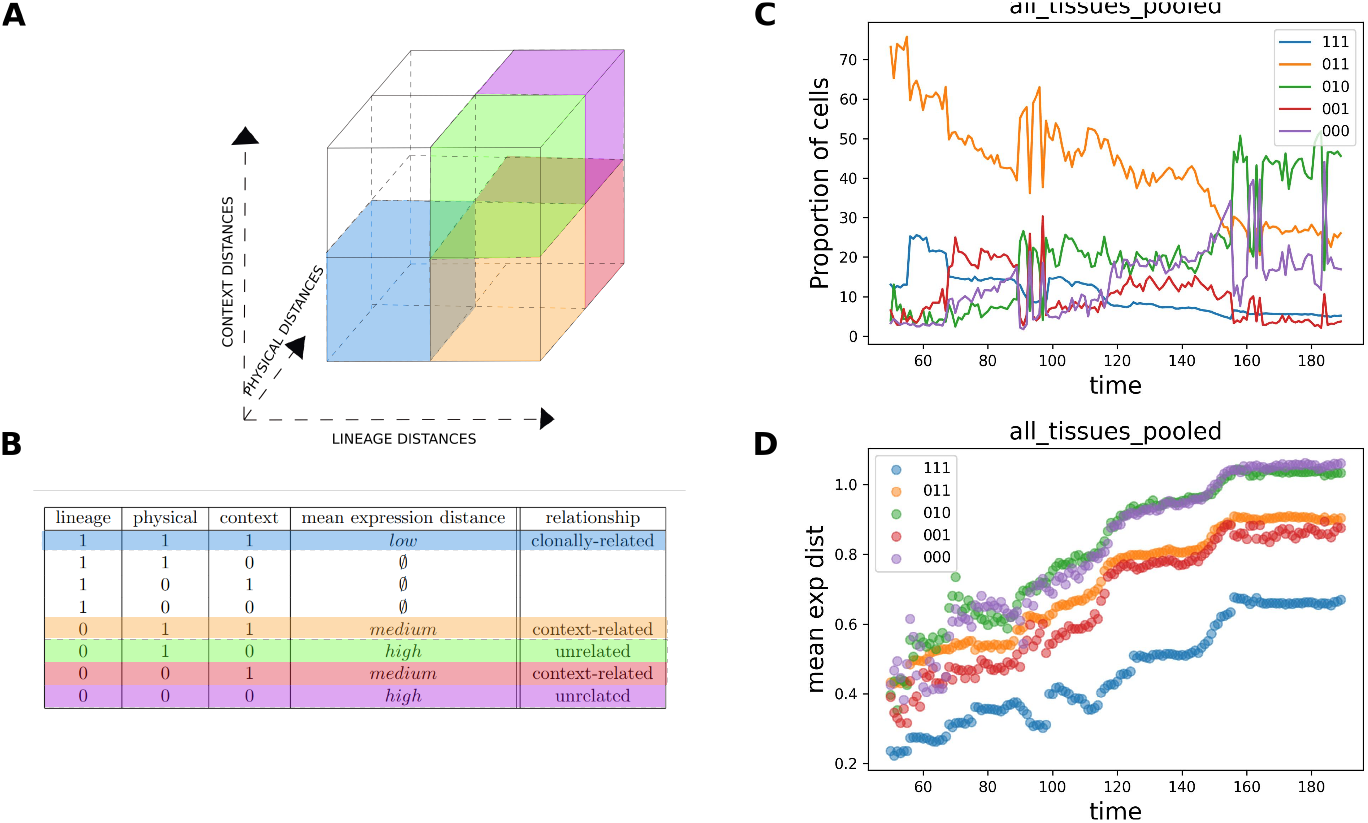
Quadrant analysis of all tissues pooled *C. elegans* A) for a given time-point, all the pairs of cells are distributed in a space according to their *lineage distance*, physical distance and *context distance*. This space is divided into 8 regions, using the mid distance between minimal and maximal values as separation lines. B) Characteristics of each configuration corresponding to the regions drawn in A). “1” represents “close” (between the minimal value and the mid-line) and “0” represents “far” (between the mid-line and the maximal value). Three of the configurations are not occupied : there are no pairs of cells that are close in lineage without being also close in context and in space. Mean *expression distance* column refers to D). C)Proportion of cells in each configuration in time D) Mean *expression distances* in each configuration in time

Moreover, for a pair of cells that are close in space, we anticipate that they share a certain proportion of their neighbors. In this case, the fact that they have a low *context distance* is entangled with the fact that they are closely related in lineage. We checked that beyond a *spatial distance* of 3 edges apart in the Delaunay graph, there are almost no shared neighbor anymore (see Supplementary Figure 2). Analyzing the mean ratio of shared neighbors between all pairs of cells in time, we see that from t=50, this ratio is sufficiently low to allow to consider that the *context distance* does not only reflect shared neighbourhood.

### Analyzing the relationship between *lineage distance* and *spatial distance* reveals that clonally-related cells are responsible for the high lineage dependency contribution

We then sought to understand better the lineage and context contributions by including in the analysis all the 4 distances measured. If we consider *lineage, physical* and *context distances*, and a binary value: close(1) or far(0), we can distribute the pairs into a total of 8 quadrants that represent all the configurations as show in Figure 3. First, we realise that some quadrants are not represented in the data: cells that are closely related in lineage, are not far in the physical space. And cells that are closely related in lineage, close in physical space are not far in terms of *context distances*. This excludes 3 quadrants. In the remaining 5 configurations, we can measure the percentage of cells in each configuration over time, as well as the mean *expression distance*. These 5 configurations represent (i) clonal cells which just only divided from a common ancestor, still close in space and in terms of context, (ii) cells that are close in context and in space, while being distant in lineage, (iii) cells that are close in context while being both distant in lineage and in space (this configuration is particularly interesting illustration of the regulative mode), (iv) border cells that are close in space without being close in lineage nor context (v) unrelated cells. The *expression distances* are low in clonal cells (i), mild in (ii) and (iii) and high in (iv) and (v). This adds two additional interpretations of the stratified correlation plots, Figure 2. The high value of lineage contribution could only be due to the low values of *expression distances* associated to clonal cells (i) when all other existing configurations (ii,iii,iv,v) that have high *lineage distance* have higher *expression distances*. In addition, the context-dependency that we observed in Figure 2 has an important contribution of cells that are close in context but not in space (prevalence of (iii) and its mild *expression distances*). This analysis by configuration can also be done for each tissue separately, which gives additional insights as shown in supplementary.

When classifying the pairs of cells in the 8 quadrants of all the possible configurations, there are three quadrants that are not occupied: in average over time between 0.3 and 0.7 % of pairs of cells belong to these quadrants. There are no pairs of cells that are close in lineage while being far spatially or/and in context. There is no occurrence of configuration where cells are close only in lineage. This is also the case in the lineage-determined model and the random null model. However the quadrant of cells that are clonally related (in blue) has very low *expression distance* with respect to the others which are not lineage-related. This adds another interpretation of the stratified correlation measure of lineage dependency contribution: this contribution is high mainly due to the contribution of cells that are clonally related.

### The changes in the modes of specification are correlated with proliferation and spatial rearrangements

In the case of the Skin specification, which was shown in Figure 2 to display a high level of context dependency compared to lineage dependency at late times, we discuss the mutual contributions of division rate and rearrangements. The results are shown on Figure 4.

**Figure 4.**
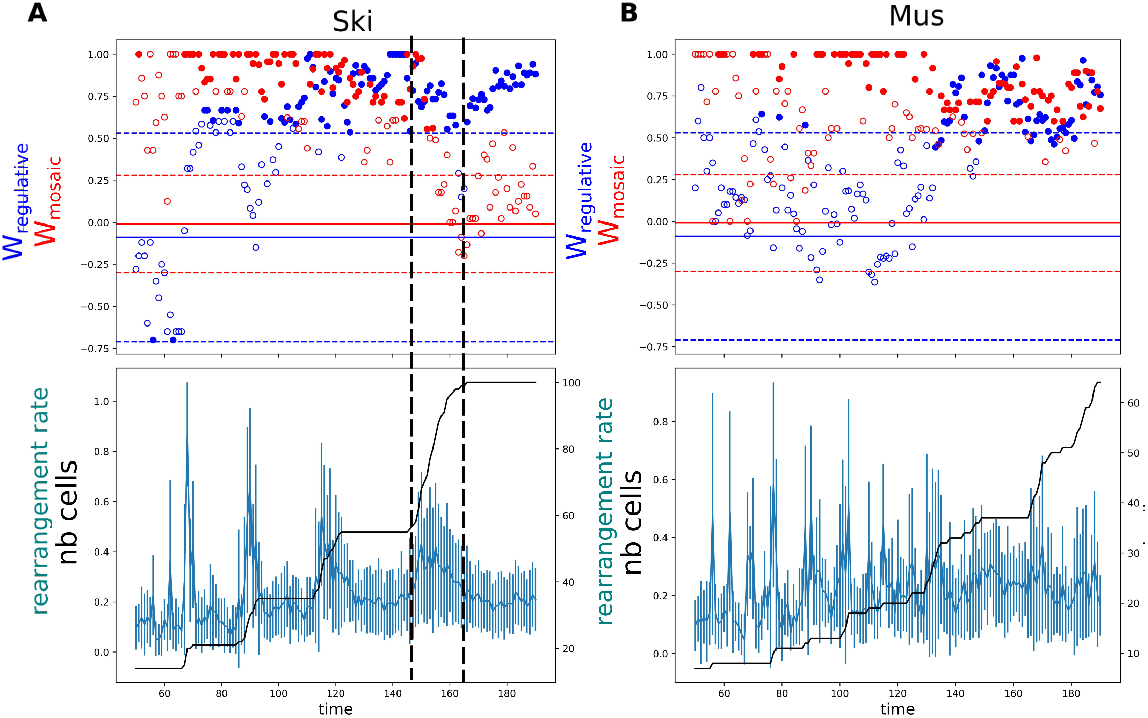
Relationship between specification and individual cell dynamics, cell proliferation and cellular spatial rearrangement. A) Top panel shows the lineage and context contributions measured by *W*_*mosaic*_ and *W*_*regulative*_ in Skin tissue. The bottom panel, with shared time point x-axis, shows the rearrangement rate in blue and the number of cells that belong to Skin cells lines in black. The vertical dotted lines delimit the last tracked division round. The daughter cells generated during this round are context determined according to our measure (significant context *W*_*regulative*_ in blue and non significant lineage *W*_*mosaic*_ in red). After this round of divisions, whereas no new cells are generated, the context *W*_*regulative*_ keeps increasing, suggesting sorting by cellular spatial rearrangement. B) For comparison, in Muscle cells, the division rounds are more asynchronous and new cells are generated until the end of the tracking

The dominant mode of specification, lineage or context determined, can be put in relation with the cell proliferation dynamics and cellular spatial rearrangement. For example, in Skin specification, there is steep drop of the lineage dependency contribution, as marked between the dashed lines in Figure 4 that coincides with a round of divisions, visible through the increase in steep slope of the number of cells leading to skin cells. The new cells that just divided have adopted expressions that correlate more with their context than with with their lineage inheritance. However, after the round of divisions is over, when the plateau of number of cells has been reached, the context dependency contribution keeps increasing, even though we only have one measure of expression state per cell cycle and there is an absence of generation of new cells with new expressions. Thus only cellular rearrangement is responsible of this increase. The rearrangement rate is measured as mean of the jaccard distances between the identities of the neighboring cells of a pair of cells. This rate is between 10% and 20%, staying in same range in all the periods without divisions. In the end, when the context dependency contribution increases whereas the number of cells stays constant, there must be targeted rearrangements, or sorting process, that place similar cells in similar contexts.

To illustrate another kind of behavior, we show the case of muscle cells. The divisions seem more asynchronous, with no such clear plateaus. The rearrangement rate stays in similar range.

### An example of genealogically distant cells converging to the same cell fate following distinct trajectories

We have shown in previous sections that there are pairs of cells which have similar context and expression, while being genealogically and physically distant: the red quadrant in Figure3. We focus on a particular pair of cells that belong to this quadrant at the last time point (t=190). These cells are named are ABplaapppp (green) and Cpappd (red). They are very distant genealogically : their lines diverge as soon as the first division, when the egg P0 divides into AB and P1, ABplaapppp deriving from AB and Cpappd from P1 (Figure 5B). We represent in Figure 5 the trajectories that lead to these two cells, which have converged to the same fate. We define “trajectories” as follows. The two cells are considered endpoint of a list constituted by the successive cells along a genealogical line. The spatial and expression trajectories are made of, respectively, the successive physical locations (Figure 5C), and positions in the high dimensional expression space (Figure 5A), of cells in this list. We show that these two trajectories converge to the same cell fate following very distinct trajectories in both physical and expression spaces. Their physical distance increases in time up to 5 edges away in spatial network of cells (Figure 5 C and F). They end up at the skin layer of the worm, in different regions, one on the posterior-right part (Cpadppd in red) and the other on the left-ventral part (ABplaapppp in green). Their *expression distance*, normalized in time, to account for the overall divergence of expression as differentiation proceeds, drops at t=155. This convergence in expression is concomitant to the drop in normalized *context distance* (Figure 5D and E) which drops progressively from t=136 to t=155.

**Figure 5.**
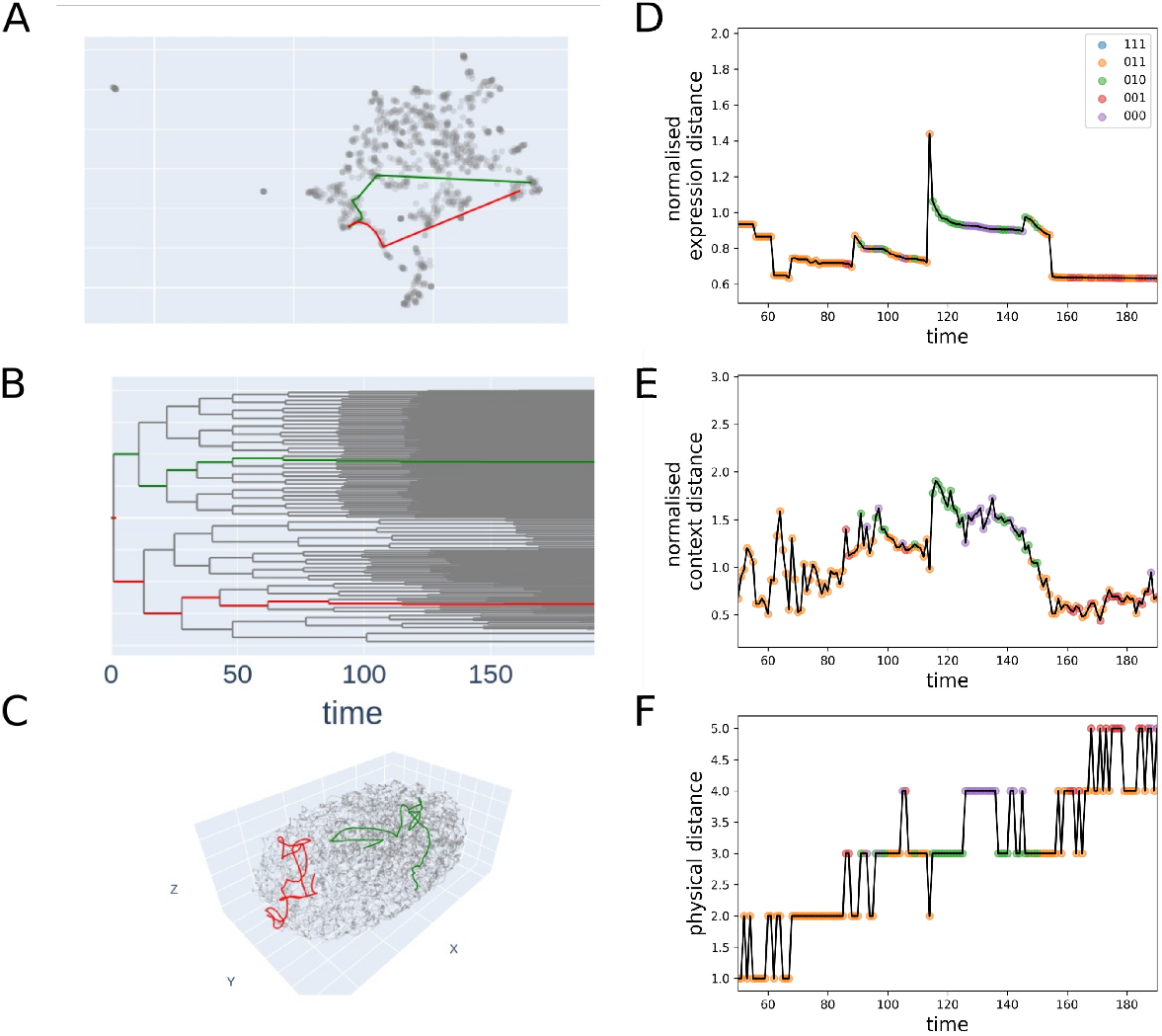
Illustration of two individual cell tracks that lead to a pair of cells that are close in expression and in context but far in lineage and in space. A) UMAP representation of the expression of all cells generated across development. The red and green lines represent the tracks that lead to the pair of cells. B) the tracks in lineage. C) the tracks in 3D space. D) *expression distance* between the two tracks normalized by the mean *expression distance* between all the pairs of cells of each time point. This confirms the convergence of cells expression visible in the UMAP in A). The colors of the dots correspond to the quadrants defined in Figure3 E) the *context distances* decrease at about the same time as the *expression distances* F) the physical distances increase. The pair of cells occupy different regions of the skin.

## Discussion

By considering jointly spatial positions, lineage information and expression at single-cell resolution, we proposed a new methodology to assess the contributions of mosaic and regulative modes of development. Looking at distances instead of the value of individual parameters directly enabled us to reduce the complexity of the problem by reducing the dimensionality and focusing on the interactions between cells. We showed using the *C. elegans* model system that those two modes could both contribute to the development at various times in the various tissues: the regulative mode is at play earlier in Neurons and Skin specification than in Pharynx and Muscle. In the range of time we are considering, it is not at play in Intestine specification. The mosaic mode is at play in all tissues and all times considered except at the end of Skin specification. We have seen that the effect of the context can be isolated in some cases, whereas the pairs of cells close in lineage are also close in space and in context so that the effects remain entangled. Then our results suggested that the causality between context and expression could be explained by cellular sorting of similar cells into similar contexts. Last, we show an illustration of a pair of cells distantly related in lineage that converge to the same cell fate through different paths in physical and expression space. This convergence is accompanied by a drop in *context distances*.

As an additional measure for the regulative contribution, it could also be interesting to consider contact areas as it was shown that they play a role in the strength of cell-cell interactions [25]. A recent paper makes a morphological atlas of *C. elegans* development at single-cell resolution available, so that even the contact area could be included, for example as weights on the edges of the spatial network [24]. Moreover, we chose to focus on protein expression as a proxy for the state of a cell. It could be interesting to go further and include their transcriptional states [4], it should be noted nevertheless that currently this comprehensive transcriptomic atlas [4] contains a significant number of transcriptomic profiles that cannot be identified at single-cell resolution due to left right symmetry of the embryo [26], restricting our ability to map each unique transcriptomic profile to a cell in *C. elegans*.

A limitation of our interpretation is that we measure the correlation between the *expression distances* and the *context distances* (respectively the *lineage distances*) in all the pairs of cells existing at the same given time point. We use this correlation as a proxy for regulative (respectively mosaic) mode. In some cases, alternative interpretation could be invoked such as in the Skin cells in the last rounds of divisions, where the high correlation between expression and *context distances* are due to cells sorting into similar neighbourhoods, as suggested earlier by Schnabel et al. [27] [28]. A clearer causal link showing the dominance of the regulative mode, would have been supported by small *context distances* preceding small *expression distances* in time.

However, the measure of such causal link is difficult. Indeed, even if the temporal resolution of the 3D positions is high enough to trace the displacements of a given cell within one cell cycle, the expression profiles are measured only once for each cell so that the dynamics of expression lacks precision [5].

Thanks to the lineage invariance of *C. elegans*, we could create directly an integrated dataset from published sources [5, 13]. The questions raised in this article are however general questions that could be addressed on other model organisms, where one can acquire and match lineage, spatial, and expression single-cell data. These types of datasets are likely to become more widespread. Recent development in high-throughput data acquisition techniques and combined to machine learning based data integration techniques have enabled the generation of large datasets using single cell RNASeq [29], lineage tracing [8], spatial transcriptomics [30], time series [31] and multi-omics datasets [32].

We have restricted our analysis to a correlation analysis, partly because of the low temporal resolution of our datasets. An increased temporal resolution in the molecular state of the cells would enable finer analysis of the processes. The computation of the Granger causality for example [33], would lead to prediction of the causal mechanisms based on time series analysis, without any perturbation of the process. Alternatively, one can imagine that modeling from biophysical mechanisms [10, 34] could be a way to assess the causality links between the variety of variables at play in the system.

Finally, previous work has analysed lineage trees in animals development in terms of algorithmic complexity [35]. In *C. elegans* in particular, it was shown that the complexity is lower than expected by chance but not as low as it could be. The main reason proposed for this sub-optimality, despite evolutionary pressure, comes from the requirement to place the adequate cells in their particular positions. We show here, that this even more true as there is a clear role for the spatial neighborhood of each cell in the determination of their fate. Our results confirm that development is a process that unfolds in space and time, involving both lineage-inherited and space-dependent factors at the single-cell level.

## Acknowledgments

We thank Vincent Bertrand, Pierre Recouvreux, Khulganaa Buyannemekh, Raphaël Clément, Charlotte Rulquin, Florence Bansept, Dominic Skinner for fruitful discussions.

---

**Supplementary Figure 1:**
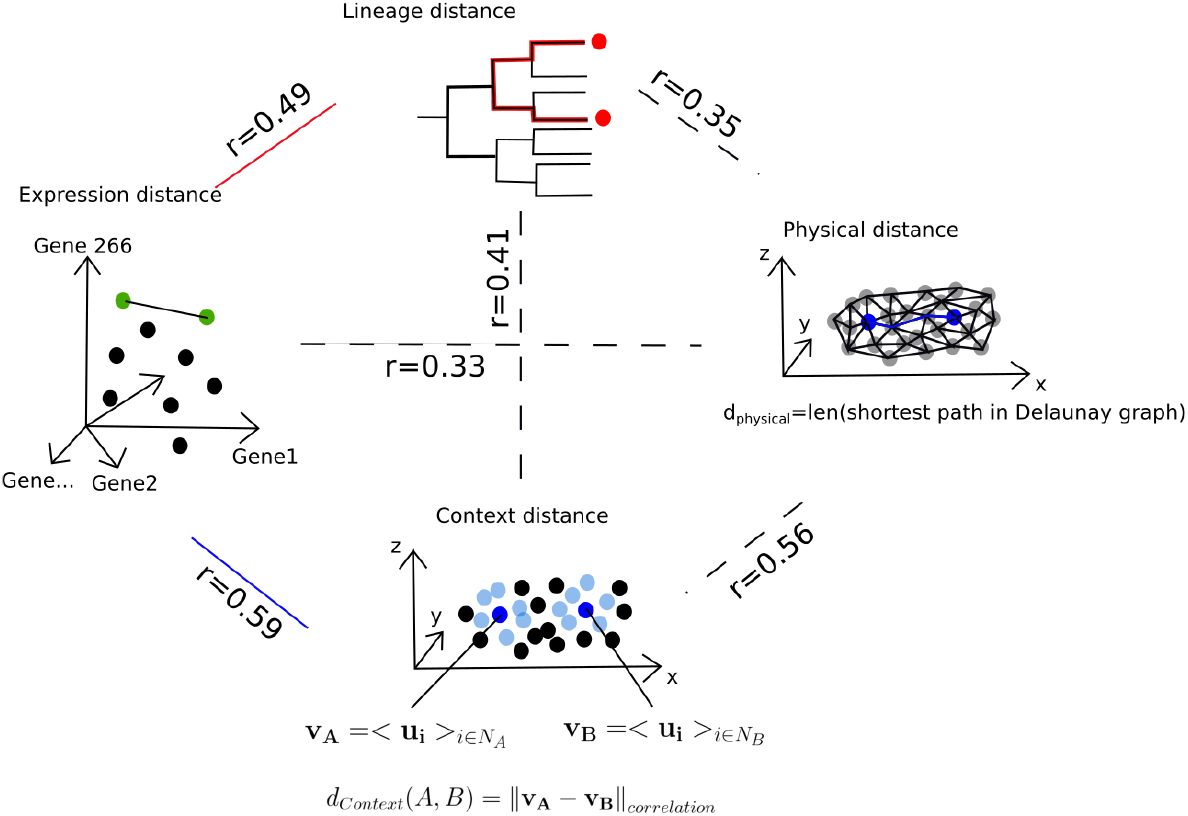
Pearson correlation r between all pairs of measured distances

**Supplementary Figure 2:**
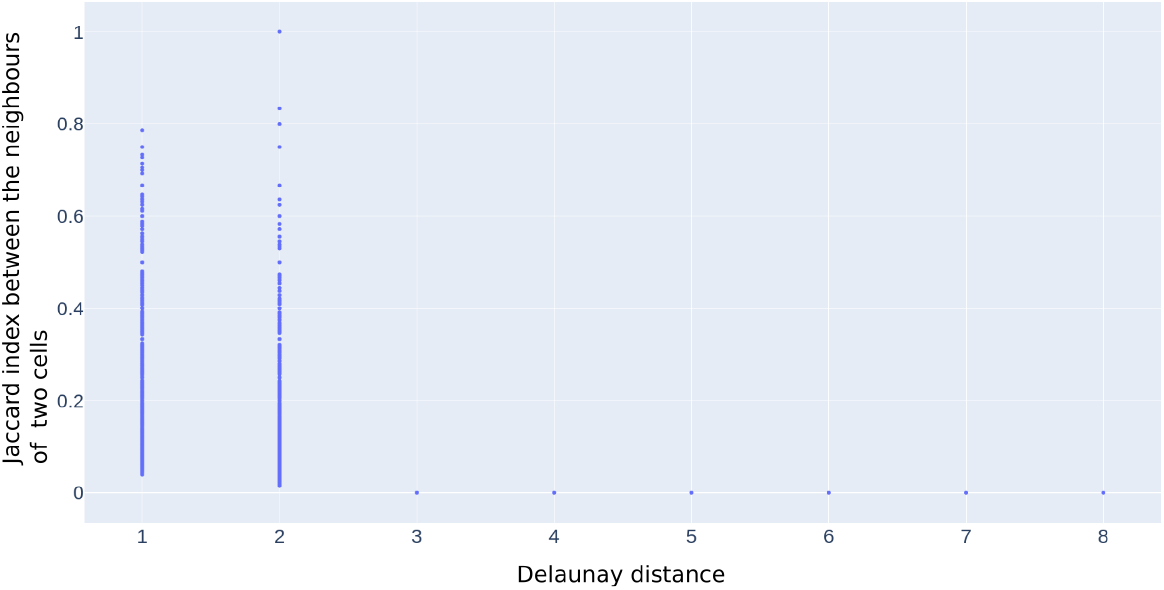
Proportion of shared neighbours between two cells, calculated as the Jaccard index between the two sets of neighbours, with respect to the physical distance between the two cells, calculated as the shortest length in the Delaunay graph

## References

1. Claudio Collinet and Thomas Lecuit. Programmed and self-organized flow of information during morphogenesis. Nature Reviews Molecular Cell Biology, 22(4):245–265, 2021.

2. Scott E Fraser and Richard M Harland. The molecular metamorphosis of experimental embryology. Cell, 100(1):41–55, 2000.

3. Noriyuki Satoh. Developmental biology of ascidians. (No Title), 1994.

4. Jonathan S Packer, Qin Zhu, Chau Huynh, Priya Sivaramakrishnan, Elicia Preston, Hannah Dueck, Derek Stefanik, Kai Tan, Cole Trapnell, Junhyong Kim, et al. A lineage-resolved molecular atlas of c. elegans embryogenesis at single-cell resolution. Science, 365(6459):eaax1971, 2019.

5. Xuehua Ma, Zhiguang Zhao, Long Xiao, Weina Xu, Yahui Kou, Yanping Zhang, Gang Wu, Yangyang Wang, and Zhuo Du. A 4d single-cell protein atlas of transcription factors delineates spatiotemporal patterning during embryogenesis. Nature Methods, 18(8):893–902, 2021.

6. Hanna L Sladitschek, Ulla-Maj Fiuza, Dinko Pavlinic, Vladimir Benes, Lars Hufnagel, and Pierre A Neveu. Morphoseq: full single-cell transcriptome dynamics up to gastrulation in a chordate. Cell, 181(4):922–935, 2020.

7. Wouter Saelens, Robrecht Cannoodt, Helena Todorov, and Yvan Saeys. A comparison of single-cell trajectory inference methods. Nature biotechnology, 37(5):547–554, 2019.

8. Daniel E Wagner and Allon M Klein. Lineage tracing meets single-cell omics: opportunities and challenges. Nature Reviews Genetics, 21(7):410–427, 2020.

9. Nicolas Olivier, Miguel A Luengo-Oroz, Louise Duloquin, Emmanuel Faure, Thierry Savy, Israël Veilleux, Xavier Solinas, Delphine Débarre, Paul Bourgine, Andrés Santos, et al. Cell lineage reconstruction of early zebrafish embryos using label-free nonlinear microscopy. Science, 329(5994):967–971, 2010.

10. Paul Villoutreix, Julien Delile, Barbara Rizzi, Louise Duloquin, Thierry Savy, Paul Bourgine, René Doursat, and Nadine Peyriéras. An integrated modelling framework from cells to organism based on a cohort of digital embryos. Scientific reports, 6(1):37438, 2016.

11. Aaron McKenna, Gregory M Findlay, James A Gagnon, Marshall S Horwitz, Alexander F Schier, and Jay Shendure. Whole-organism lineage tracing by combinatorial and cumulative genome editing. Science, 353(6298):aaf7907, 2016.

12. Duncan M Chadly, Kirsten L Frieda, Chen Gui, Leslie Klock, Martin Tran, Margaret Y Sui, Yodai Takei, Remco Bouckaert, Carlos Lois, Long Cai, et al. Reconstructing cell histories in space with image-readable base editor recording. bioRxiv, pages 2024–01, 2024.

13. Xiaoyu Li, Zhiguang Zhao, Weina Xu, Rong Fan, Long Xiao, Xuehua Ma, and Zhuo Du. Systems properties and spatiotemporal regulation of cell position variability during embryogenesis. Cell reports, 26(2):313–321, 2019.

14. Ralf Schnabel. Why does a nematode have an invariant cell lineage? In Seminars in cell & developmental biology, volume 8, pages 341–349. Elsevier, 1997.

15. JE Sulston and JG White. Regulation and cell autonomy during postembryonic development of caenorhabditis elegans. Developmental biology, 78(2):577–597, 1980.

16. Stephan W Grill, Pierre GoÉnczy, Ernst HK Stelzer, and Anthony A Hyman. Polarity controls forces governing asymmetric spindle positioning in the caenorhabditis elegans embryo. Nature, 409(6820):630–633, 2001.

17. Nathan W Goehring, Philipp Khuc Trong, Justin S Bois, Debanjan Chowdhury, Ernesto M Nicola, Anthony A Hyman, and Stephan W Grill. Polarization of par proteins by advective triggering of a pattern-forming system. Science, 334(6059):1137–1141, 2011.

18. Bijan Etemad-Moghadam, Su Guo, and Kenneth J Kemphues. Asymmetrically distributed par-3 protein contributes to cell polarity and spindle alignment in early c. elegans embryos. Cell, 83(5):743–752, 1995.

19. Ivan PG Moskowitz and Joel H Rothman. lin-12 and glp-1 are required zygotically for early embryonic cellular interactions and are regulated by maternal glp-1 signaling in caenorhabditis elegans. Development, 122(12):4105–4117, 1996.

20. Vincent Bertrand, Paul Bisso, Richard J Poole, and Oliver Hobert. Notch-dependent induction of left/right asymmetry in c. elegans interneurons and motoneurons. Current Biology, 21(14):1225–1231, 2011.

21. Giovanni Lo Iacono, Alasdair JC Cook, Gianne Derks, Lora E Fleming, Nigel French, Emma L Gillingham, Laura C Gonzalez Villeta, Clare Heaviside, Roberto M La Ragione, Giovanni Leonardi, et al. A mathematical, classical stratification modeling approach to disentangling the impact of weather on infectious diseases: A case study using spatio-temporally disaggregated campylobacter surveillance data for england and wales. PLoS computational biology, 20(1):e1011714, 2024.

22. Naomi Altman and Martin Krzywinski. Points of significance: Association, correlation and causation. Nature methods, 12(10), 2015.

23. Nikolaos Papadopoulos, Parra R Gonzalo, and Johannes Söding. Prosstt: probabilistic simulation of single-cell rna-seq data for complex differentiation processes. Bioinformatics, 35(18):3517–3519, 2019.

24. Jianfeng Cao, Guoye Guan, Vincy Wing Sze Ho, Ming-Kin Wong, Lu-Yan Chan, Chao Tang, Zhongying Zhao, and Hong Yan. Establishment of a morphological atlas of the caenorhabditis elegans embryo using deep-learning-based 4d segmentation. Nature communications, 11(1):6254, 2020.

25. Léo Guignard, Ulla-Maj Fiúza, Bruno Leggio, Julien Laussu, Emmanuel Faure, Gaël Michelin, Kilian Biasuz, Lars Hufnagel, Grégoire Malandain, Christophe Godin, et al. Contact area–dependent cell communication and the morphological invariance of ascidian embryogenesis. Science, 369(6500):eaar5663, 2020.

26. Malek Senoussi, Thierry Artieres, and Paul Villoutreix. Partial label learning for automated classification of single-cell transcriptomic profiles. PLOS Computational Biology, 20(4):e1012006, 2024.

27. Ralf Schnabel, Marcus Bischoff, Arend Hintze, Anja-Kristina Schulz, Andreas Hejnol, Hans Meinhardt, and Harald Hutter. Global cell sorting in the c. elegans embryo defines a new mechanism for pattern formation. Developmental biology, 294(2):418–431, 2006.

28. Marcus Bischoff and Ralf Schnabel. Global cell sorting is mediated by local cell–cell interactions in the c. elegans embryo. Developmental biology, 294(2):432–444, 2006.

29. Chengxiang Qiu, Beth K Martin, Ian C Welsh, Riza M Daza, Truc-Mai Le, Xingfan Huang, Eva K Nichols, Megan L Taylor, Olivia Fulton, Diana R O’Day, et al. A single-cell time-lapse of mouse prenatal development from gastrula to birth. Nature, pages 1–10, 2024.

30. Anjali Rao, Dalia Barkley, Gustavo S França, and Itai Yanai. Exploring tissue architecture using spatial transcriptomics. Nature, 596(7871):211–220, 2021.

31. Mingyue Wang, Qinan Hu, Zhencheng Tu, Lingshi Kong, Jiajun Yao, Rong Xiang, Zhan Chen, Yan Zhao, Yanfei Zhou, Tengxiang Yu, et al. A single-cell 3d spatiotemporal multi-omics atlas from drosophila embryogenesis to metamorphosis. bioRxiv, pages 2024–02, 2024.

32. Zhen He, Shuofeng Hu, Yaowen Chen, Sijing An, Jiahao Zhou, Runyan Liu, Junfeng Shi, Jing Wang, Guohua Dong, Jinhui Shi, et al. Mosaic integration and knowledge transfer of single-cell multimodal data with midas. Nature Biotechnology, pages 1–12, 2024.

33. Jungsik Noh, Tadamoto Isogai, Joseph Chi, Kushal Bhatt, and Gaudenz Danuser. Granger-causal inference of the lamellipodial actin regulator hierarchy by live cell imaging without perturbation. Cell systems, 13(6):471–487, 2022.

34. Julien Delile, Matthieu Herrmann, Nadine Peyriéras, and René Doursat. A cell-based computational model of early embryogenesis coupling mechanical behaviour and gene regulation. Nature communications, 8(1):13929, 2017.

35. Ricardo BR Azevedo, Rolf Lohaus, Volker Braun, Markus Gumbel, Muralikrishna Umamaheshwar, Paul-Michael Agapow, Wouter Houthoofd, Ute Platzer, Gaëtan Borgonie, Hans-Peter Meinzer, et al. The simplicity of metazoan cell lineages. Nature, 433(7022):152–156, 2005.

